# Structural and functional significance of arginine at position 2 of angiotensin II: Implications for oxidant-mediated amino acid residue conversions

**DOI:** 10.1101/2025.06.28.662142

**Authors:** Yuichiro J. Suzuki, Yukako C. Tamazawa, Fariha E. Bablu, Emily S. Gonzales, Trwska S. Ghafoor, Chani S. Chung, Alessia M. Suzuki, Amber R. Mickelson, Eliza J. Murphy, Jon Hao, Tadahisa Teramoto

## Abstract

Angiotensin II (Ang II) is a major mediator of pathophysiological processes. Angiotensin-converting enzyme 2 (ACE2) regulates Ang II levels by cleaving it into Ang 1-7. Our previous study showed that arginine at the second position (Arg2) of Ang II plays an important role in regulating ACE2 peptidase activity. The present study examined the mechanisms and functions of Arg2 in Ang II. Cleavage of Arg2 from Ang II, or single amino acid replacement of a positively charged Arg with other amino acids, particularly negatively charged amino acids such as glutamic acid (Glu), eliminated Ang II’s ability to compete with the fluorogenic substrate used in the ACE2 activity assay. Arg2-to-Glu substitution also affected the ability of angiotensin I and angiotensin III to interact with ACE2. These results confirmed the important roles of Arg2 in the ACE2 enzymatic mechanism. Since Arg and proline (Pro) residues are susceptible to protein carbonylation, and both residues produce glutamic semialdehyde, which can be further oxidized to Glu, we previously proposed that Arg, Pro, and Glu could be interchangeable, and that ROS promote amino acid conversions within protein structures post-translationally in biological systems. Thus, Arg2 of Ang II could undergo oxidant-mediated amino acid residue conversion to become Glu or Pro. The presence of both circulating Ang II (Arg2Glu) and Ang II (Arg2Pro) in rats were confirmed by mass spectrometry. Global gene expression analyses revealed that Arg2Glu- and Arg2Pro-converted Ang II peptides elicit different cellular signaling responses compared with wild-type Ang II, suggesting that Arg2-modifed Ang II peptides exhibit distinct biological functions.

## Introduction

Angiotensin II (Ang II), a peptide hormone, is an essential regulator of vascular tone and blood pressure. In addition to its physiologically important functions, this peptide also mediates various pathological conditions [Forrester et al., 2018; Paz Ocaranza et al., 2020]. In the renin-angiotensin pathway, angiotensinogen, a protein synthesized in the liver, is cleaved by renin in the kidney to form angiotensin I (Ang I), a 10-amino acid peptide with the sequence Asp-Arg-Val-Tyr-Ile-His-Pro-Phe-His-Leu. Ang I then travels through the circulation to the lung, where angiotensin-converting enzyme (ACE) cleaves it to generate Ang II, an 8-amino-acid-peptide (Asp-Arg-Val-Tyr-Ile-His-Pro-Phe) [Forrester et al., 2018; Paz Ocaranza et al., 2020; Benigni et al., 2010; Yang et al., 2011]. The major actions of Ang II are to bind to the angiotensin II type 1 receptor and promote various pathophysiological processes, including smooth muscle contraction, smooth muscle cell growth, aldosterone and antidiuretic hormone production, sympathetic nervous system tone, cardiac hypertrophy, and fibrosis [Paz Ocaranza et al., 2020; Yang et al., 2011].

The recent pandemic of Coronavirus Disease 2019 (COVID-19) heightened awareness of the importance of another angiotensin-regulatory enzyme, angiotensin-converting enzyme 2 (ACE2), which was identified as the major host cell receptor for the COVID-19 virus, Severe Acute Respiratory Syndrome Coronavirus 2 (SARS-CoV-2), for infection [Gheblawi et al., 2020]. Physiologically, ACE2 cleaves *C*-terminal Phe of Ang II to produce Ang 1-7 (Asp-Arg-Val-Tyr-Ile-His-Pro). This cleavage of the results in the conversion of the vasoconstrictor Ang II into the vasodilator Ang 1-7 [Saulnier et al., 2024]. Thus, ACE2 plays a critical role in regulating Ang II levels and functions under both physiological and pathological conditions.

In addition for Ang II to bind to its receptors, Ang II can be cleaved at the *N*-terminus by aminopeptidases to form Ang III (Arg-Val-Tyr-Ile-His-Pro-Phe), which can then be further cleaved to Ang IV (Val-Tyr-Ile-His-Pro-Phe) [Padia et al., 2008]. While Ang III functions similarly to Ang II by binding to Ang II receptors (i.e AT1 or AT2 receptors), Ang IV mainly binds to a distinct receptor (the AT4 receptor), which plays a role in brain signaling and memory [De Bundel et al., 2008].

In addition to Ang II to producing these shorter peptides, we recently proposed that Ang II oxidation by reactive oxygen species (ROS) converts amino acid residues in angiotensin peptides [Bablu et al., 2026]. This hypothesis is based on the rationale that oxidation of both Arg and Pro produces glutamic semialdehyde, which contains a carbonyl group [Berlett & Stadtman, 1997; Amici et al., 1989], and that glutamic semialdehyde can be further oxidized to Glu; thus, these 3 amino acid residues could be interchangeable [Suzuki & Hao, 2017; Suzuki, 2019]. This concept of oxidant-mediated amino acid residue conversions could be applied to the Ang II molecule [Bablu et al., 2026], as this peptide possesses Arg at the second position (Arg2), which was previously shown to be important for Ang II to serve as a substrate for ACE2 [Saulnier et al., 2024]. We found that both Ang II and Ang III effectively competed with a fluorogenic ACE2 substrate in an ACE2 enzymatic assay; this ability was lost upon cleavage of Arg2 to produce Ang IV [Saulnier et al., 2024].

The present study further characterized the role of Arg2 in Ang II in ACE2 peptidase activity. As we provide mechanistic insights into the relationships between this Arg and angiotensin peptides, we also obtained important results identifying, in the biological system, the novel angiotensin peptides in which Arg2 is modified.

## Materials and Methods

### ACE2 Activity Assay

The Fluorimetric SensoLyte 390 ACE2 Activity Assay Kit (AnaSpec Inc., Fremont, CA, USA; Catalog # AS-72086) was used to monitor ACE2 carboxylase activity using a FRET-based artificial substrate. Cleavage of the Mca/Dnp FRET peptide (Mca-Ala-Pro-Lys-Dnp; 25 µM) to the fluorescent Mca (7-Methoxycoumarin-4-acetic acid) peptide catalyzed by recombinant human ACE2 (40 pM; Sino Biological, Inc., Wayne, PA, USA) in the presence or absence of angiotensin peptides was monitored using the POLARstar OPTIMA Microplate Reader (BMG Labtech, Cary, NC, USA) at Ex/Em = 320 nm/420 nm. The MCA reference standard curve generated by our system indicated that 1,000 relative fluorescent units (RFU) correspond to 0.3 µM product. Experiments we performed showed linear fluorescence-concentration responses. Angiotensin peptides were synthesized by contracting with Aapptec LLC (Louisville, KY) at ≥ 95% purity (confirmed by HPLC and mass spectrometry) and used at 40 µM unless otherwise noted.

### Molecular Dynamics Simulation

The ACE2 structure (PDB ID: 1R42) was obtained from the Protein Data Bank and prepared by removing non-essential ligands while retaining the Zn²□ ion. Ang II (DRVYIHPF) and its mutants [R2P (DPVYIHPF), R2E (DEVYIHPF), R2K (DKVYIHPF), and R2Q (DQVYIHPF)], as well as Ang I (DRVYIHPFHL), Ang III (RVYIHPF), Ang IV (VYIHPF), and Ang (1–7) (DRVYIHP), were constructed using molecular modeling tools on the Neurosnap AI platform (https://neurosnap.ai) and then underwent energy minimization. The peptides were docked into the ACE2 active site (Zn core) using the ClusPro 2.0 protein–protein docking server (https://cluspro.org), which provided coefficient weights for evaluating the docking results [Kozakov et al., 2017]. The server also provided three-dimensional models of the docked complexes, enabling visualization and validation of ligand conformations within the ACE2 binding pocket. The final refined complex structures were visualized using PyMOL (PyMOL Molecular Graphics System, Version 2.4.1, Schrödinger, LLC). The Zn core distance was calculated in PyMOL, and the docking models were generated with SMINA using the AutoVina docking tool.

### Nanospray LC/MS/MS analysis

Serum samples collected from Fischer 344 testing rats were analyzed in this study. In brief, the short peptides were extracted by using 60% Acetonitrile and purified by C18 column. The purified peptides mixtures from each sample was reconstituted in 0.1% formic acid before nanospray liquid chromatography/mass spectrometry/mass spectrometry (LC/MS/MS) analysis was performed. The peptides mixture from each sample was analyzed using a Thermo Scientific Exploris 480 Orbitrap Mass Spectrometer (Thermo Electron, Bremen, Germany) equipped with a Thermo Vanquish Neo RSLCnano System (Thermo Electron, Bremen, Germany). Peptide samples were loaded onto a peptide trap cartridge at a flow rate of 5 μl/min. The trapped peptides were eluted onto a reversed-phase Easy-Spray Column PepMap RSLC, C18, 2 µM, 100A, 75 µm × 250 mm (Thermo Scientific) using a linear gradient of acetonitrile (3-36%) in 0.1% formic acid. The elution duration was 60 min at a flow rate of 0.3 μl/min. Eluted peptides from the Easy-Spray column were ionized and sprayed into the mass spectrometer, using a Nano Easy-Spray Ion Source (Thermo) under the following settings: spray voltage, 1.6 kV, Capillary temperature, 275°C. The instrument was operated in the data-dependent mode to automatically switch between full scan MS and MS/MS acquisition. Survey full scan MS spectra (m/z 300−2,000) were acquired in the Orbitrap with 70,000 resolution (m/z 200) after the accumulation of ions to a 3 × 106 target value based on predictive AGC from the previous full scan. Dynamic exclusion was set to 20 s. The 15 most intense multiply charged ions (z ≥ 2) were sequentially isolated and fragmented in the HCD cell using normalized HCD collision energy at 28% with an AGC target of 1e5 and a maximum injection time of 100 ms at 17,500 resolution. The raw MS files were analyzed using the Thermo Proteome Discoverer 3.1 (Thermo Scientific, Bremen, Germany) for peptide identification. Raw data files were searched against the target angiotensin peptide sequences database using the Proteome Discoverer 3.1 software (Thermo, San Jose, CA) based on the SEQUEST algorithm. The minimum peptide length was specified to be five amino acids. The precursor mass tolerance was set to 15 ppm, whereas fragment mass tolerance was set to 0.01 Da. The maximum false peptide discovery rate was specified as 0.05. The resulting Proteome Discoverer Report contains all assembled peptides sequences and peptide spectrum match counts (PSM#) and MS1 peak area.

### Cell culture

Human pulmonary vascular smooth muscle cells (Catalog # 3110) were purchased from ScienCell Research Laboratories (Carlsbad, CA, USA). The cells were cultured according to the manufacturer’s instructions at 37°C in 5% CO_2_. Cells at passage four were treated in triplicate with Ang II (wild-type), Ang II (Arg2Glu), or Ang II (Arg2Pro) peptides dissolved in deionized water (dH_2_O). A final concentration of 100 nM Ang II peptides was used, consistent with previous publications [Griendling et al., 1994; Takahashi et al., 1999; Eguchi et al., 2001; Leung et al., 2013]. An equal volume of dH_2_O was added to the negative controls. Three hours after cell treatment, total RNA was extracted using TRIzol (Thermo Fisher Scientific, Waltham, MA, USA) and quantified using a 2100 Bioanalyzer (Agilent Technologies, Santa Clara, CA, USA).

### RNA-Seq transcriptome analysis

The Illumina paired-end mRNA-Seq library construction and RNA-seq sequencing were performed by Azenta Life Sciences (South Plainfield, NJ). Using the Zymo-Seq RiboFree Total RNA Library Kit (Zymo Research, Irvine, CA, USA), each RNA sample was purified and reverse-transcribed to cDNAs, which was then ligated with the P7 adaptor sequence at the 3’ end, followed by second-strand synthesis and P5 adaptor ligation at the opposite sites of the double-stranded DNA. After purification based on DNA size (300-600bp) using the beads provided in the kit, index PCR was performed. Successful libraries were confirmed with the D1000 ScreenTape Assay (Agilent Technologies) on a TapeStation and sequenced on an Illumina Novaseq to a depth of >30 million read pairs (150 bp paired-end sequencing). Individual paired read data (R1 and R2) were collected in separate fastq files.

Read mapping, transcript assembly, and differential gene expression analyses were performed using Geneious Prime (Biomatters, v. 2024.7.0, Auckland, New Zealand), as previously used in gene expression analysis studies [Samaras et al., 2020; Espinosa et al., 2019; Hoermann et al., 2022]. Sequence reads in fastq files (R1 and R2) were set as a paired-end and mapped to the *Homo Sapiens* genome (https://www.ncbi.nlm.nih.gov) using the “Map to Reference” function. Transcript counts were obtained using the “Calculate Expression Levels” function. Normalized TPM (transcripts per million) counts were generated to account for differences in the total numbers of mRNAs. Differential expression log_2_ fold-change ratios were determined using the “Compare Expression Levels” function. Differentially expressed genes were identified as having an absolute value of log_2_ fold-change greater than 1 or less than –1 and a *p*-value less than 0.05. Volcano plots were generated using the log_2_ fold change value and *p*- value on the SRplot website (https://www.bioinformatics.com.cn/srplot) [Tang et al., 2023].

*Statistical Analysis:*

Means and standard errors of the mean (SEM) were computed. Two groups were compared using a two-tailed Student’s t-test, and differences among more than two groups were assessed using the analysis of variance (ANOVA). P < 0.05 was considered statistically significant.

## Results

### Role of Arg2 of Ang II in interactions with ACE2

Ang II can be cleaved by aminopeptidases at the *N*-terminus to produce Ang III and Ang IV. Fig. 1A shows the amino acid sequences of Ang II, Ang III, and Ang IV. The fluorescence resonance energy transfer (FRET)-based ACE2 peptidase activity assay was used to investigate the ability of Ang II and other angiotensin peptides to compete with the fluorogenic ACE2 substrate (Mca-Ala-Pro-Lys-Dnp). In this system, cleavage of the Pro-Lys linkage by ACE2 eliminates quenching of fluorescence signals via FRET, thereby enabling monitoring of ACE2 peptidase activity. As expected, Ang II (a natural substrate for ACE2) effectively competed with the fluorogenic ACE2 substrate (Fig. 1B) in a dose-dependent fashion (Fig. 1C). Ang III, in which *N*-terminal aspartic acid (Asp) is deleted from Ang II, also competed with the fluorogenic substrate for ACE2 (Figs. 1B and 1C). By contrast, the truncation of arginine (Arg) at the 2^nd^ position (Arg2) of Ang II significantly reduced the ability to compete with the fluorogenic ACE2 substrate (Figs. 1B and 1C). Consistently to these experimental results, molecular dynamics simulation revealed that Ang II and Ang III, but not Ang IV, enter the ACE2 active site, so that *C*-terminal Phe of angiotensin peptides becomes proximal to the active site zinc ion (Fig. 1D). Molecular modeling calculations further showed that Ang IV possesses less favorable docking coefficient weight (i.e. less estimated binding affinity) toward ACE2 compared to Ang II or Ang III (Fig. 1E). These results together indicate the importance of Arg2 of the angiotensin peptides for Ang II and Ang III correctly interacting with the ACE2 enzyme as substrates. ACE2 activity assay experiments in which both Ang II and Ang IV were added together showed that the effects of Ang II + Ang IV were like Ang II alone (Fig. 1F), indicating that Ang II enters the ACE2 active site efficiently.

**Fig. 1:**
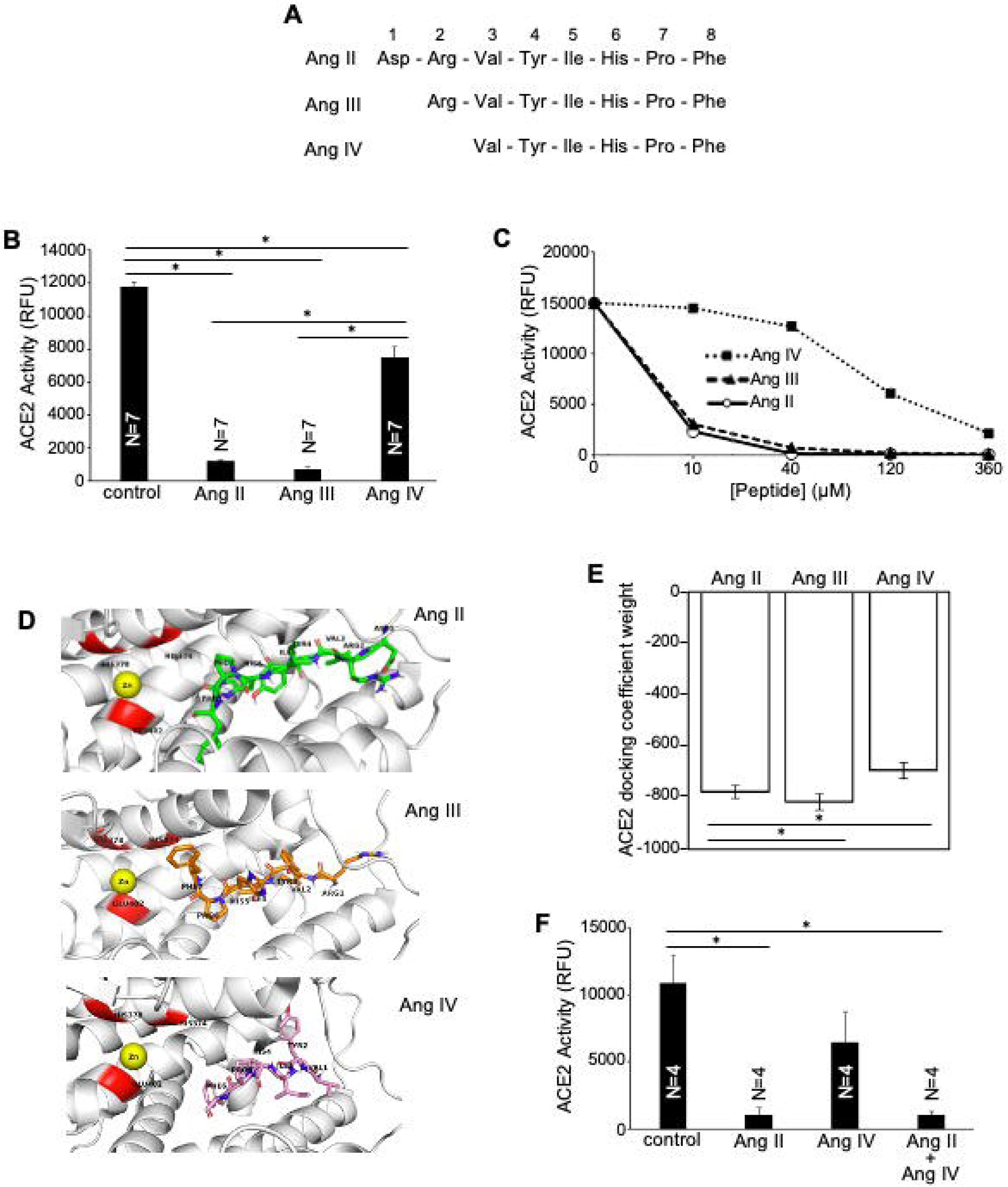
Effects of *N*-terminal truncation of Ang II on functional and structural interactions with ACE2. (A) Amino acid sequences of Ang II, Ang III, and Ang IV. (B) Effects of Ang II, Ang III, and Ang IV on ACE2 activity assay. ACE2, a FRET-based fluorogenic substrate (Mca-Ala-Pro-Lys-Dnp) at 25 µM, and Ang II, Ang III, or Ang IV at 40 µM were incubated and the ACE2 peptidase activity was monitored. Results show that Ang II and Ang III, but not Ang IV, almost completely inhibited the cleavage of the fluorogenic ACE2 substrate. (C) Dose-response of the effects of Ang II, Ang III, and Ang IV on ACE2 activity. (D) Molecular dynamics simulations were employed to model the characteristics of Ang II, Ang III, and Ang IV within the ACE2 substrate tunnel. The simulation indicates that Ang II and Ang III enter the ACE2 substrate tunnel with the *C*-terminal Phe located proximal to the zinc (Zn) catalytic site, while the *C*-terminal Phe of Ang IV is located less proximal. (E) Calculated ACE2 docking coefficient weight values, where more negative values reflect stronger, more favorable binding affinity, indicating that Ang IV has less binding affinity toward the ACE2 active site compared to Ang II or Ang III. (F) ACE2 activity assay results showing that Ang IV does not interfere with Ang II to compete with the fluorogenic substrate. The symbol (*) denotes values that are different from each other at *p* < 0.05.

To further confirm the functional role of Arg2 in the Ang II peptide, we replaced Arg at the position 2 with other amino acids. Replacing Arg with the simpler amino acid Ala (Arg2Ala) significantly reduced Ang II’s ability to compete with the fluorogenic substrate for ACEs, although Ang II (Arg2Ala) still exhibited some competitive ability (Figs. 2A and 2B). Ang II in which Arg2 is replaced by Lys, another positively charged amino acid (Arg2Lys), had minimal effect on Ang II’s ability to compete with the fluorogenic ACE2 substrate (Figs. 2A and 2B). On the other hand, replacing Arg2 (a positively charged amino acid) in Ang II with Glu or Asp (negatively charged amino acids) almost eliminated Ang II’s ability to compete with the fluorogenic ACE2 substrate, whereas uncharged Gln did not (Fig. 2C-E). Molecular modeling predicted that Ang II (Arg2Glu) exhibits a less favorable docking coefficient weight than wild-type Ang II or Ang II (Arg2Lys), suggesting weaker binding to ACE2 (Fig. 2F).

**Fig. 2:**
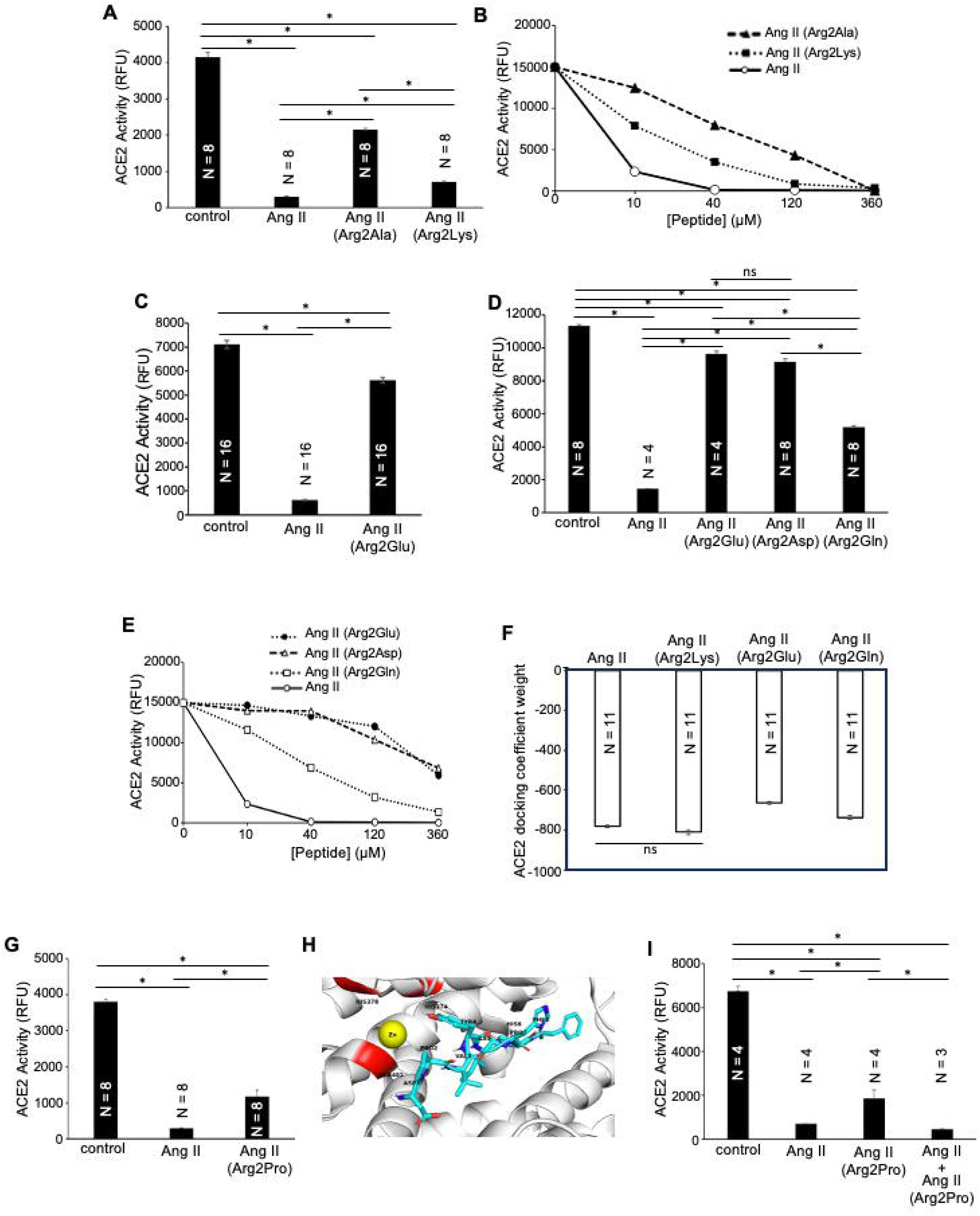
Effects of replacing Arg2 of Ang II with other amino acids on functional and structural interactions with ACE2. (A) Effects of Ang II, Ang II (Arg2Ala), and Ang II (Arg2Lys) on ACE2 activity assay. Results show that replacing Arg2 with Ala significantly inhibited Ang II’s ability to compete with the fluorogenic substrate, whereas replacing the positively charged Arg with another positively charged Lys had minimal effect. (B) Dose-response of the effects of Ang II, Ang II (Arg2Ala), and Ang II (Arg2Lys) on ACE2 activity. (C) Replacing the positively charged Arg with the negatively charged Glu almost eliminated Ang II’s ability to compete with the fluorogenic substrate. (D) Similarly, replacing the positively charged Arg to a negatively charged Asp, but not neutral Gln, almost eliminated the ability of Ang II to compete with the fluorogenic substrate. (E) Dose-response of the effects of Ang II, Ang II (Arg2Glu), Ang II (Arg2Asp) and Ang II (Arg2Gln) on ACE2 activity. (F) Calculated ACE2 docking coefficient weight values, where more negative values reflecting stronger, more favorable binding affinity, estimating that Ang II (Arg2Glu) has the least binding affinity toward the ACE2 active site, whereas that of Ang II (Arg2Lys) is comparable to Ang II. (G) Effects of replacing Arg2 of Ang II with Pro on ACE2 activity. (H) Molecular dynamics simulation depicting the characteristics of Ang II (Arg2Pro) in the ACE2 substrate tunnel. The simulation indicates that, in contrast to Ang II, Ang II (Arg2Pro) enters the ACE2 substrate tunnel with the *N*-terminus proximal to the catalytic site. (I) ACE2 activity assay results showing that Ang II (Arg2Pro) does not interfere with Ang II to compete with the fluorogenic substrate. The symbol (*) denotes values that are different from each other at *p* < 0.05.

Replacing Arg2 in Ang II with Pro produced intermediate effects, i.e., less efficient than Ang II but still had some effects (Fig. 2G). Molecular dynamics simulations revealed that, while wild-type Ang II enters the ACE2 zinc-binding pocket with its *C*-terminal Phe positioned near the catalytic site (Fig. 1D), Ang II (Arg2Pro) approaches the enzyme from the *N*-terminal side (Fig. 2H). ACE2 activity assay experiments in which both Ang II and Ang II (Arg2Pro) were added showed that the effects of these two peptides together were similar to those of Ang II alone (Fig. 2I), indicating that Ang II enters the ACE2 active site efficiently.

### Role of N-terminal Arg of Ang III in interactions with ACE2

We next examined the Arg residue of Ang III using peptides in which this amino acid is replaced with Glu or Pro (Fig. 3A). As *N*-terminal Asp is deleted from Ang II, Ang III possesses Arg as the *N*-terminal amino acid. As described in Fig. 1B, Ang III exhibits similar inhibitory effects to Ang II on ACE2 activity as assessed using the fluorogenic substrate, suggesting that Ang II and Ang III similarly enter the ACE2 active site. Replacing the positively charged Arg residue of Ang III with a negatively charged Glu dramatically attenuated Ang III’s ability to compete with the fluorogenic substrate, whereas replacing Arg to Pro had minimal effect (Figs. 3B and 3C). Similarly to Figs. 1F and 2I, strong inhibitory effects of Ang II on ACE2 activity, competing with the fluorogenic substrate, persisted even in the presence of fewer effective peptides (Figs. 3D and 3E).

**Fig. 3:**
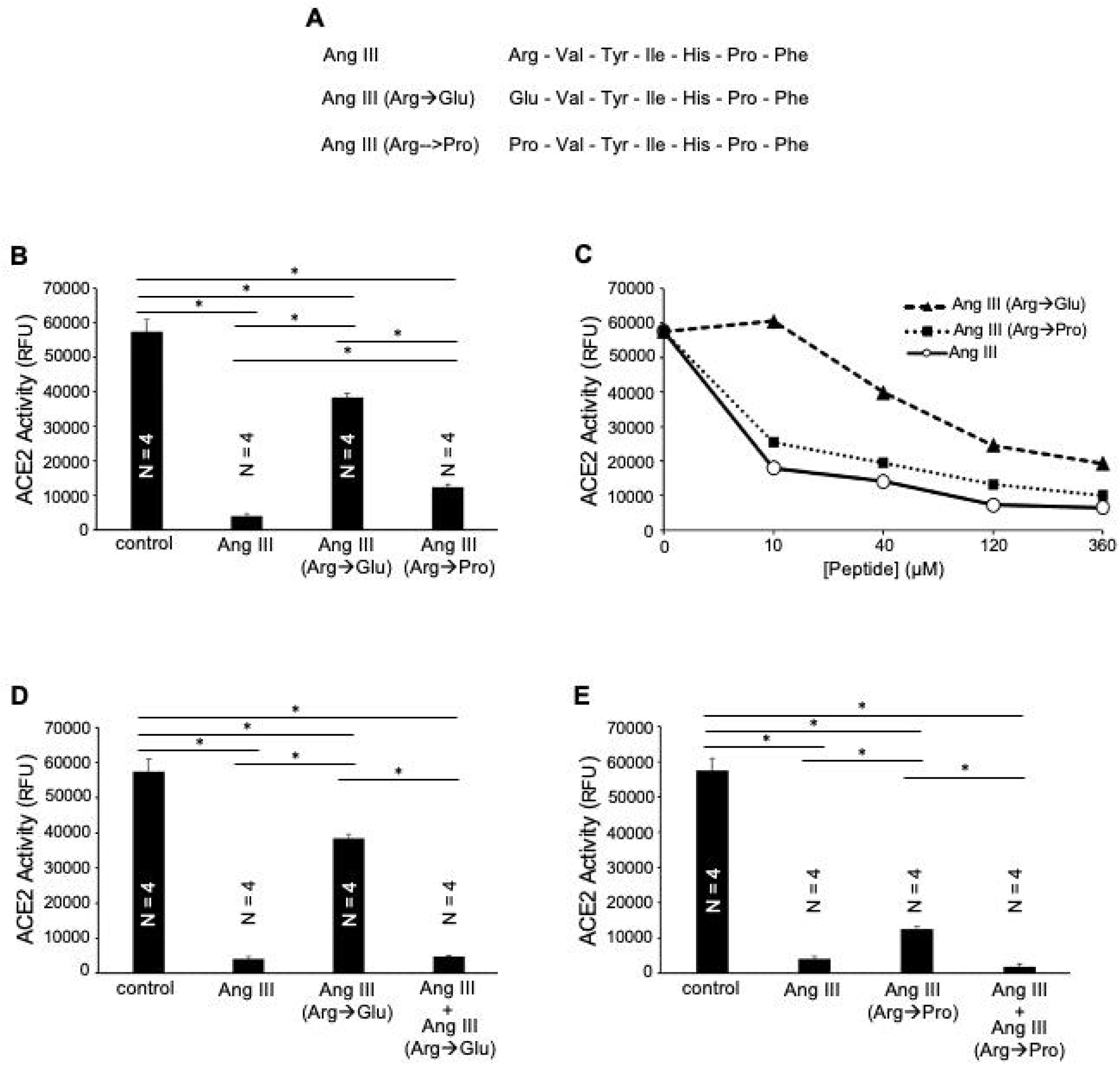
Effects of replacing Arg of Ang III with Glu or Arg on ACE2 activity. (A) Amino acid sequences of Ang III, Ang III with Arg replaced with Glu (Arg➜Glu), and Ang III with Arg replaced with Pro (Arg➜Pro). (B) Effects of Ang III, Ang III (Arg➜Glu), and Ang III (Arg➜Pro) on ACE2 activity assay. (C) Dose-response. (D) ACE2 activity assay results showing that Ang III (Arg➜Glu) does not interfere with Ang III to compete with the fluorogenic substrate. (E) ACE2 activity assay results showing that Ang III (Arg➜Pro) does not interfere with Ang III to compete with the fluorogenic substrate. The symbol (*) denotes values that are different from each other at *p* < 0.05.

### Role of Arg2 of Ang I in interactions with ACE2

Ang I also serves as a substrate for ACE2 in a reaction to produce Ang 1-9 [Donoghue et al., 2000]. However, ACE2 activity assays showed that Ang I is less effective than Ang II at competing with the fluorogenic ACE2 substrate (Fig. 4A). Consistent with this, molecular dynamics simulations estimated that the *C*-terminal Phe of Ang II docks closer to the catalytic Zn²□ ion (2.3 Å) than the *C*-terminal Leu of Ang I (4.1 Å), as visualized in molecular modeling of Ang II (Fig. 1D) and Ang I (Fig. 4D) fitting into the ACE2 catalytic site. As described above for Ang II and Ang III, replacing the positively charged Arg residue at the second position of Ang I with a negatively charged Glu inhibited Ang I’s ability to compete with the fluorogenic substrate for ACE2, whereas Pro substitution had lesser effects (Figs. 4E and 4F).

**Fig. 4:**
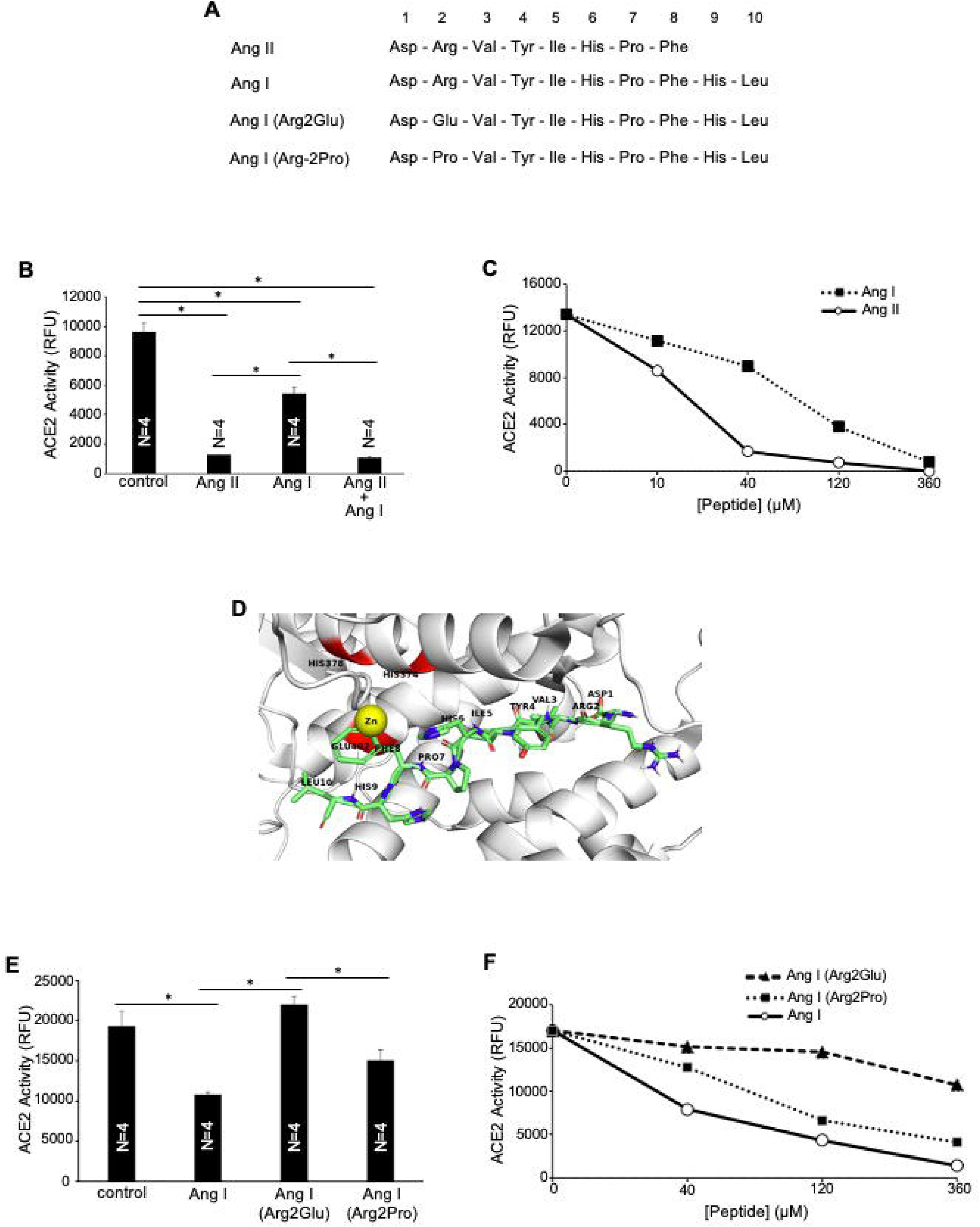
Effects of Ang I on ACE2 activity. (A) Amino acid sequences of Ang II, Ang I, Ang I (Arg2Glu), and Ang I (Arg2Pro). (B) Effects of Ang II, Ang I, and a combination of Ang I and Ang II on the ACE2 activity assay, showing that Ang I competes with the fluorogenic substrate with less efficacy than Ang II and that Ang I does not interfere with the ability of Ang II. (C) Dose-response of the effects of Ang II and Ang I on ACE2 activity. (D) Molecular dynamics simulation showing that Ang I enters the ACE2 substrate tunnel with the *C*-terminal side located proximal to the catalytic site. (E) Effects of Ang I, Ang I (Arg2Glu), and Ang I (Arg2Pro) on ACE2 activity. (F) Dose response. The symbol (*) denotes values that are different from each other at *p* < 0.05.

### Effects of Ang 1-7 on ACE2 activity

ACE2 cleaves Ang II to produce Ang 1-7 [Jiang et al., 2014]. Molecular dynamics simulations showed that, as an ACE2 product, Ang 1-7 can enter the substrate tunnel but remains distal to the zinc active site (Fig. 5A) and has an ACE2 docking coefficient weight (Fig. 5B) like Ang IV or Ang II (Arg2Glu) that exhibited minimal abilities to compete with the fluorogenic substrate. Nevertheless, Ang 1-7 showed some ability to compete with the fluorogenic substrate for ACE2 (Figs. 5C and 5D). As with other angiotensin peptides described above, replacing the negatively charged Arg at the second position with a positively charged Glu, Ang 1-7 (Arg2Glu), eliminated the ability of Ang 1-7 to interfere with the fluorogenic substrate (Figs. 5C and 5D). By contrast to other angiotensin peptides, the Pro substitution at Arg2 in Ang 1-7 (Arg2Pro) also resulted in complete inhibition of competition with the fluorogenic substrate (Figs. 5C and 5D). The ability of Ang 1-7 to somewhat interfere with the fluorogenic substrate remained even in the presence of Ang 1-7 (Arg2Glu) or Ang 1-7 (Arg2Pro).

**Fig. 5:**
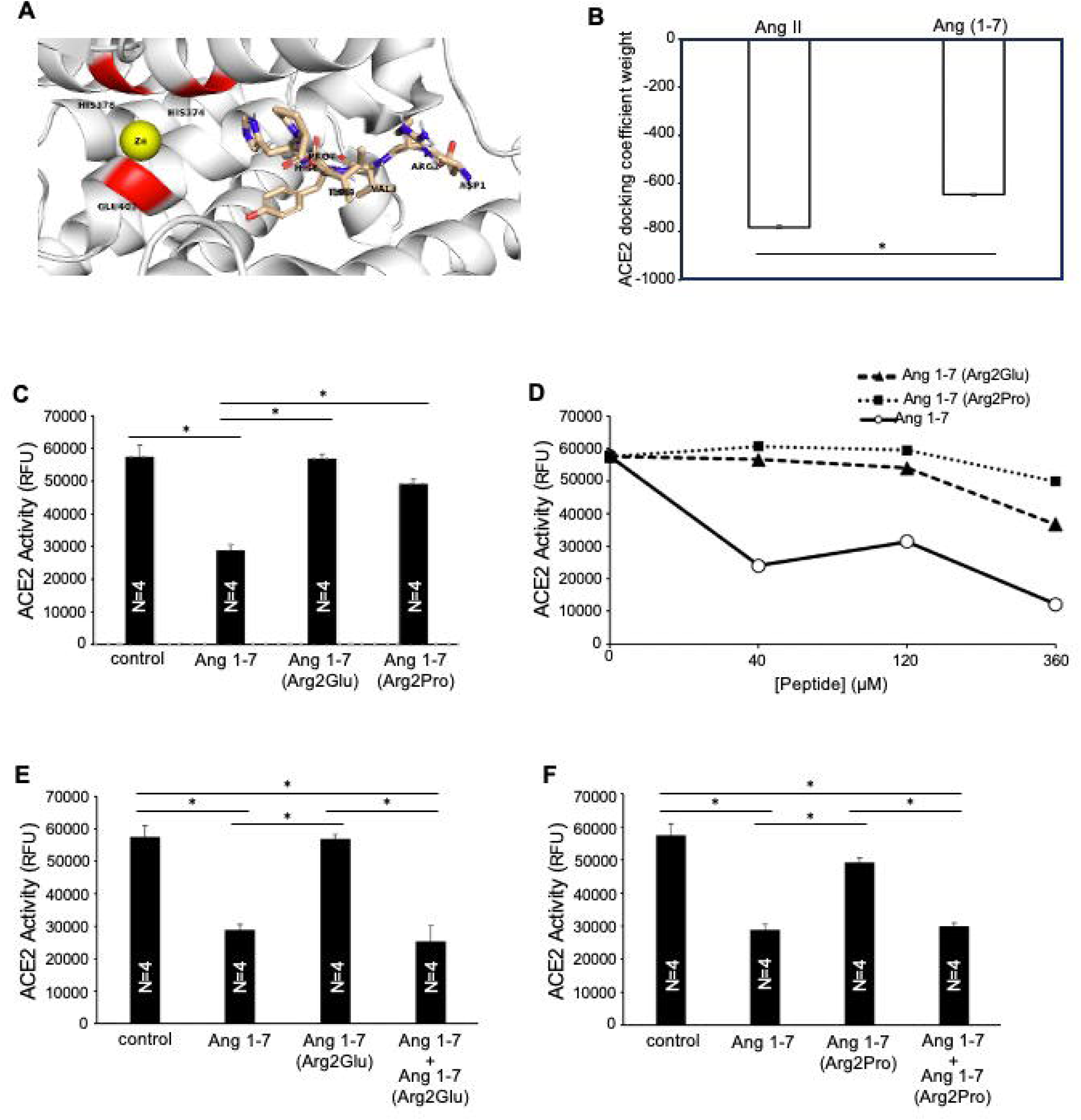
Effects of Ang 1-7 on ACE2 activity. (A) Molecular dynamics simulation showing the characteristics of Ang 1-7 in the ACE2 substrate tunnel. (B) Calculated ACE2 docking coefficient weight values, estimating that Ang 1-7 has less binding affinity toward the ACE2 active site than Ang II. (C) Effects of Ang 1-7, Ang 1-7 (Arg2Glu), and Ang 1-7 (Arg2Pro) on ACE2 activity. (D) Dose-response. (E) Effects of Ang 1-7, Ang 1-7 (Arg2Glu), and a combination of Ang 1-7and Ang 1-7 (Arg2Glu) on ACE2 activity assay. (F) Effects of Ang 1-7, Ang 1-7 (Arg2Pro), and a combination of Ang 1-7and Ang 1-7 (Arg2Pro) on ACE2 activity assay. The symbol (*) denotes values that are different from each other at *p* < 0.05.

### Identification of naturally-occurring Arg2-modified angiotensin peptides

We previously proposed the oxidant-mediated amino acid residue conversion process, in which Arg, Pro, and Glu residues in protein or peptide structures can be converted interchangeably through glutamic semialdehyde [Suzuki & Hao, 2017; Suzuki, 2019], thus Arg2 of Ang II could be converted to Glu or Pro in the biological system [Bablu et al., 2026]. To test the hypothesis that Arg2-modified angiotensin peptides occur naturally in the biological system, serum samples from Fischer 344 rats were processed for the extraction of peptides, followed by C18 column purification and analysis by LC/MS/MS. Results revealed that these modified Ang II peptides at Arg2 are indeed naturally circulating in the blood. We found 0.05 – 0.08 pmol/ml of Ang II (Arg2Glu) and 0.04 – 0.09 pmol/ml of Ang II are circulating in the blood of normal rats. The values for Ang II (Arg2Glu) increased to 0.13 - 0.20 pmol/ml in rats treated with SARS-CoV-2 spike protein that exhibit oxidative stress [Brink et al., 2026].

In addition to modified Ang II, our mass spectrometry analysis detected the occurrence of circulating Ang I (Arg2Glu), Ang I (Arg2Pro), and Ang III (Arg1Glu) in the blood, as shown in Table 1.

**Table 1.** List of angiotensin peptides, in which Arg (at the second position of Ang II) is replaced with either Glu or Pro (consistent with oxidant-mediated amino acid residue conversion), that were identified to occur naturally in Fischer (CDF) rat blood.

| Amino acid sequence | Name of peptide |
| --- | --- |
| DEVYIHPFHL | Ang I (Arg2Glu) |
| DPVYIHPFHL | Ang I (Arg2Pro) |
| DEVYIHPF | Ang II (Arg2Glu) |
| DPVYIHPF | Ang II (Arg2Pro) |
| EVYIHPF | Ang III (Arg1Glu) |

### Effects of Arg2-modified Ang II on gene expression

To determine whether Arg2-modified Ang II peptides exhibit different biological functions from wild-type Ang II, cultured human pulmonary artery smooth muscle cells were treated with deionized H_2_O (dH_2_O; used as a vehicle control), Ang II (wild-type), Ang II (Arg2Glu), or Ang II (Arg2Pro) for gene expression analysis. Total RNA was extracted from the cells, and the cellular transcriptomes were analyzed by RNA-Seq. Differentially expressed genes were defined as those with an absolute value of log_2_ fold-change greater than 1 or less than –1. A log_2_ fold change of 1 corresponds to a 2-fold change.

Our experiments using human pulmonary artery smooth muscle cells treated for 3 hours with wild-type Ang II identified 45 genes whose mRNAs were upregulated relative to the dH_2_O-treated control (Fig. 6A). These Ang II-upregulated genes are listed in Table 2 in order of effectiveness. In this system, we did not detect any genes that were downregulated in response to wild-type Ang II.

**Fig. 6:**
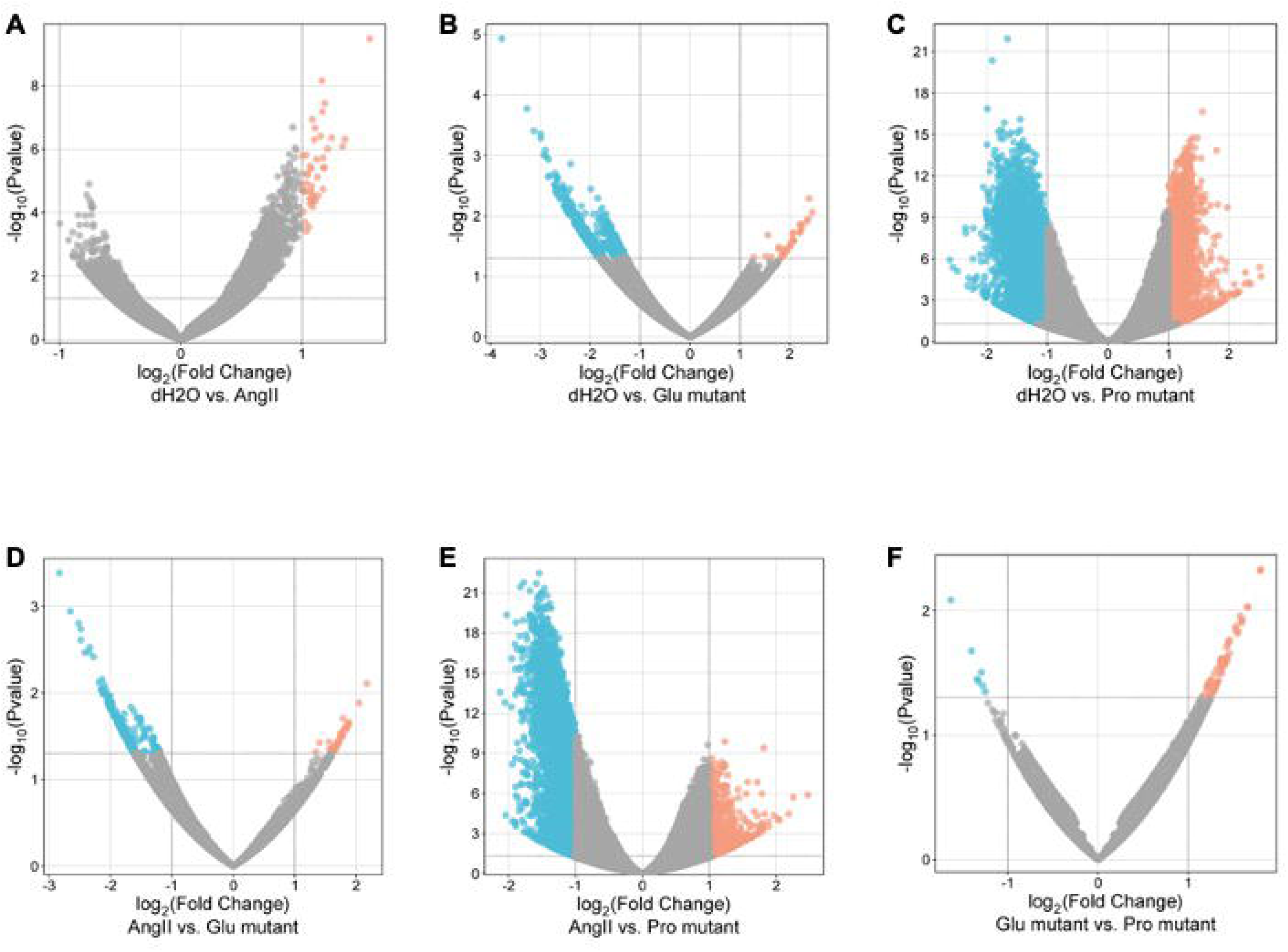
Volcano plot visualizing the relationship between the fold change and *p*-values of RNA Seq data. (A) Ang II (wild-type) relative to dH_2_O control. (B) Ang II (Arg2Glu) relative to dH_2_O control. (C) Ang II (Arg2Pro) relative to dH_2_O control. (D) Ang II (Arg2Glu) relative to Ang II (wild-type). (E) Ang II (Arg2Pro) relative to Ang II (wild-type). (F) Ang II (Arg2Pro) relative to Ang II (Arg2Glu).

**Table 2.**
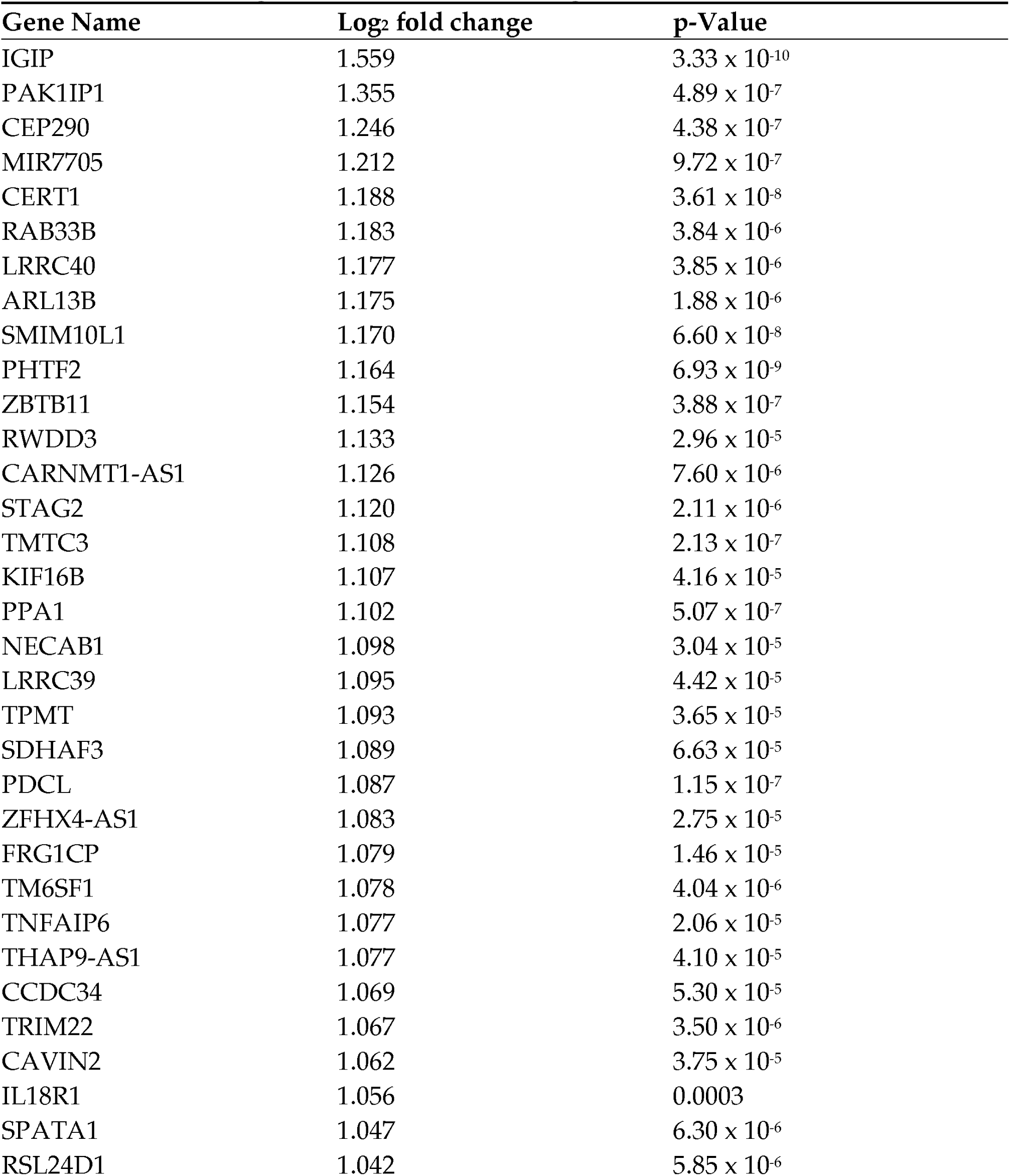
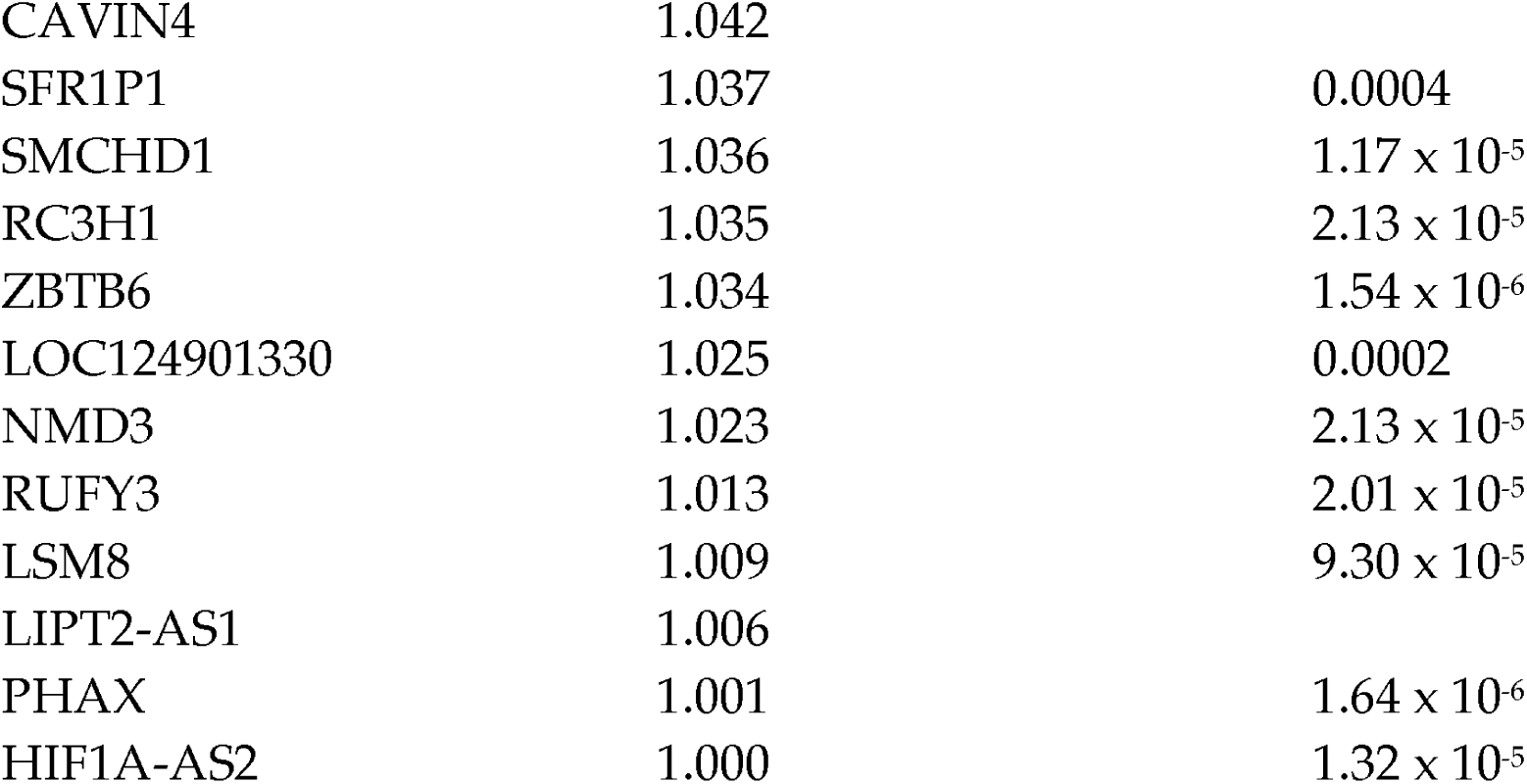
Genes upregulated in response to Ang II treatment of cells relative to dH_2_O control.

| Gene Name | Log<sub>2</sub> fold change | p-Value |
| --- | --- | --- |
| IGIP | 1.559 | $3.33 \times 10^{-10}$ |
| PAK1IP1 | 1.355 | $4.89 \times 10^{-7}$ |
| CEP290 | 1.246 | $4.38 \times 10^{-7}$ |
| MIR7705 | 1.212 | $9.72 \times 10^{-7}$ |
| CERT1 | 1.188 | $3.61 \times 10^{-8}$ |
| RAB33B | 1.183 | $3.84 \times 10^{-6}$ |
| LRRC40 | 1.177 | $3.85 \times 10^{-6}$ |
| ARL13B | 1.175 | $1.88 \times 10^{-6}$ |
| SMIM10L1 | 1.170 | $6.60 \times 10^{-8}$ |
| PHTF2 | 1.164 | $6.93 \times 10^{-9}$ |
| ZBTB11 | 1.154 | $3.88 \times 10^{-7}$ |
| RWDD3 | 1.133 | $2.96 \times 10^{-5}$ |
| CARNMT1-AS1 | 1.126 | $7.60 \times 10^{-6}$ |
| STAG2 | 1.120 | $2.11 \times 10^{-6}$ |
| TMTC3 | 1.108 | $2.13 \times 10^{-7}$ |
| KIF16B | 1.107 | $4.16 \times 10^{-5}$ |
| PPA1 | 1.102 | $5.07 \times 10^{-7}$ |
| NECAB1 | 1.098 | $3.04 \times 10^{-5}$ |
| LRRC39 | 1.095 | $4.42 \times 10^{-5}$ |
| TPMT | 1.093 | $3.65 \times 10^{-5}$ |
| SDHAF3 | 1.089 | $6.63 \times 10^{-5}$ |
| PDCL | 1.087 | $1.15 \times 10^{-7}$ |
| ZFHX4-AS1 | 1.083 | $2.75 \times 10^{-5}$ |
| FRG1CP | 1.079 | $1.46 \times 10^{-5}$ |
| TM6SF1 | 1.078 | $4.04 \times 10^{-6}$ |
| TNFAIP6 | 1.077 | $2.06 \times 10^{-5}$ |
| THAP9-AS1 | 1.077 | $4.10 \times 10^{-5}$ |
| CCDC34 | 1.069 | $5.30 \times 10^{-5}$ |
| TRIM22 | 1.067 | $3.50 \times 10^{-6}$ |
| CAVIN2 | 1.062 | $3.75 \times 10^{-5}$ |
| IL18R1 | 1.056 | 0.0003 |
| SPATA1 | 1.047 | $6.30 \times 10^{-6}$ |
| RSL24D1 | 1.042 | $5.85 \times 10^{-6}$ |
| CAVIN4 | 1.042 |  |
| SFR1P1 | 1.037 | 0.0004 |
| SMCHD1 | 1.036 | $1.17 \times 10^{-5}$ |
| RC3H1 | 1.035 | $2.13 \times 10^{-5}$ |
| ZBTB6 | 1.034 | $1.54 \times 10^{-6}$ |
| LOC124901330 | 1.025 | 0.0002 |
| NMD3 | 1.023 | $2.13 \times 10^{-5}$ |
| RUFY3 | 1.013 | $2.01 \times 10^{-5}$ |
| LSM8 | 1.009 | $9.30 \times 10^{-5}$ |
| LIPT2-AS1 | 1.006 |  |
| PHAX | 1.001 | $1.64 \times 10^{-6}$ |
| HIF1A-AS2 | 1.000 | $1.32 \times 10^{-5}$ |

To determine whether Ang II (Arg2Glu), in which Arg at position 2 is replaced with Glu, also elicits cell signaling that modifies gene expression, the same cell type was treated with Ang II (Arg2Glu) under the same conditions as those used for Ang II (wild-type). We found that Ang II (Arg2Glu) was more effective at altering gene expression, upregulating 37 genes and downregulating 674 genes (Fig. 6B). The volcano plot comparing differential gene expression in Ang II-treated cells vs. Ang II (Arg2Glu)-treated cells is also shown in Fig. 6D. Table 3 lists the genes upregulated by Ang II (Arg2Glu). Among the 45 genes that were upregulated by wild-type Ang II (Table 2), Ang II (Arg2Glu) upregulated only 2 genes: TNFAIP6 and IL18R1. These results suggest that replacing the positively charged Arg with the negatively charged Glu may block many of the actions of wild-type Ang II. While Ang II (Arg2Glu) did not upregulate most of the wild-type Ang II-responsive genes, it did upregulate other genes, including fibroblast growth factor 10 (FGF10), which increased 5-fold (calculated from log_2_ fold change of 2.330) (Table 3).

**Table 3.** Genes upregulated in response to Ang II (Arg2Glu) treatment of cells relative to dH_2_O <u>control</u>

| Gene Name | Log<sub>2</sub> fold change | p-Value |
| --- | --- | --- |
| LINC01473 | 2.448 | 0.0086 |
| LOC124901330 | 2.386 | 0.0051 |
| LOC105378145 | 2.366 | 0.0111 |
| FGF10 | 2.330 | 0.0124 |
| LOC107986764 | 2.240 | 0.0163 |
| LINC03016 | 2.223 | 0.0140 |
| LOC105379183 | 2.217 | 0.0147 |
| C2orf66 | 2.200 | 0.0137 |
| LOC105377275 | 2.199 | 0.0183 |
| LOC124900685 | 2.196 | 0.0185 |
| LOC102723724 | 2.082 | 0.0254 |
| LOC105374873 | 2.076 | 0.0257 |
| LOC105371651 | 2.072 | 0.0190 |
| DCHS2 | 2.049 | 0.0200 |
| MIOS-DT | 2.043 | 0.0253 |
| LOC124900854 | 2.012 | 0.0294 |
| LOC124901112 | 2.004 | 0.0312 |
| RPL39P18 | 1.978 | 0.0332 |
| LOC101929201 | 1.921 | 0.0375 |
| LOC124901756 | 1.914 | 0.0368 |
| RBMXP1 | 1.892 | 0.0412 |
| LOC105379080 | 1.891 | 0.0412 |
| FAM177B | 1.885 | 0.0421 |
| HSD17B13 | 1.877 | 0.0354 |
| LINC02726 | 1.874 | 0.0411 |
| LOC124909361 | 1.853 | 0.0457 |
| LOC105374226 | 1.845 | 0.0461 |
| IL26 | 1.840 | 0.0446 |
| TTPA | 1.837 | 0.0481 |
| RRH | 1.831 | 0.0482 |
| ANXA10 | 1.814 | 0.0498 |
| LOC101927888 | 1.780 | 0.0350 |
| IL18R1 | 1.774 | 0.0330 |
| EPCAM | 1.628 | 0.0476 |
| TNFAIP6 | 1.561 | 0.0206 |
| LOC124906176 | 1.520 | 0.0460 |
| SNX4 | 1.274 | 0.0481 |

In contrast to the lack of downregulation by wild-type Ang II in our experimental system, Ang II (Arg2Glu) downregulated a substantial number of genes (674 genes). A 13.6-fold decrease in the expression of *SLCO4A1* mRNA (calculated from log_2_ fold change of 3.769) was caused by Ang II (Arg2Glu). Table 4 lists twenty genes that were downregulated by Ang II (Arg2Glu), in the order of effectiveness. Tables 5 and 6 list the names of the top genes (in order of the extent of changes) that were differentially modulated (upregulated and downregulated, respectively) by Ang II (Arg2Glu) relative to Ang II (wild-type). Thus, wild-type Ang II and Ang (Arg2Glu) differentially modulate gene expression (Fig. 6D), indicating that the oxidant-mediated Arg-to-Glu conversion alters Ang II function.

**Table 4.** Genes downregulated in response to Ang II (Arg2Glu) treatment of cells relative to dH_2_O control. Top 20 genes with highest fold change are listed.

| Gene Name | Log<sub>2</sub> fold change | p-Value |
| --- | --- | --- |
| SLCO4A1 | -3.769 | 1.17 × 10<sup>-5</sup> |
| TINCR | -3.126 | 0.0004 |
| GPT | -2.992 | 0.0005 |
| CCDC116 | -2.844 | 0.0022 |
| RND1 | -2.800 | 0.0022 |
| SLC34A3 | -2.711 | 0.0030 |
| PCSK4 | -2.701 | 0.0020 |
| WDR97 | -2.686 | 0.0034 |
| MIR4758 | -2.672 | 0.0023 |
| MIP | -2.645 | 0.0045 |
| PDE6G | -2.575 | 0.0035 |
| DKFZP434A062 | -2.570 | 0.0050 |
| LHFPL4 | -2.564 | 0.0036 |
| BCAR1P1 | -2.559 | 0.0048 |
| LINC02846 | -2.548 | 0.0031 |
| BRSK2 | -2.544 | 0.0036 |
| SERTAD3-AS1 | -2.541 | 0.0040 |
| MIR6847 | -2.539 | 0.0035 |
| FREY1 | -2.522 | 0.0068 |
| SRMS | -2.510 | 0.0067 |

**Table 5.** Genes upregulated in response to Ang II (Arg2Glu) treatment of cells relative to Ang II <u>(wild-type). Seventeen genes were found to be upregulated.</u>

| Gene Name | Log<sub>2</sub> fold change | p-Value |
| --- | --- | --- |
| LINC03016 | 2.173 | 0.0078 |
| DEPDC1-AS1 | 1.837 | 0.0263 |
| MYH16 | 1.813 | 0.0264 |
| C2orf66 | 1.811 | 0.0251 |
| TLL1 | 1.785 | 0.0197 |
| HNRNPA1P59 | 1.776 | 0.0318 |
| NAIPP3 | 1.757 | 0.0287 |
| TLR5 | 1.746 | 0.0320 |
| DIO1 | 1.740 | 0.0300 |
| CFAP90 | 1.705 | 0.0392 |
| LINC02497 | 1.693 | 0.0372 |
| ANXA10 | 1.674 | 0.0392 |
| OSBPL10-AS1 | 1.671 | 0.0417 |
| SLC36A3 | 1.660 | 0.0401 |
| LINC01590 | 1.635 | 0.0417 |
| ARHGAP42P1 | 1.561 | 0.0443 |
| ADGRB3-DT | 1.400 | 0.0375 |

**Table 6.** Genes downregulated in response to Ang II (Arg2Glu) treatment of cells relative to Ang <u>II (wild-type). Top 20 genes with highest fold change are listed.</u>

| Gene Name | Log<sub>2</sub> fold change | p-Value |
| --- | --- | --- |
| SLCO4A1 | -2.833 | 0.0004 |
| FREY1 | -2.654 | 0.0011 |
| MIR10400 | -2.484 | 0.0018 |
| USHBP1 | -2.484 | 0.0024 |
| PIN1-DT | -2.419 | 0.0034 |
| BCAR1P2 | -2.350 | 0.0033 |
| MSLN | -2.337 | 0.0030 |
| MIR6847 | -2.279 | 0.0038 |
| IGSF9 | -2.189 | 0.0075 |
| CD83 | -2.156 | 0.0091 |
| PDZD4 | -2.151 | 0.0090 |
| EFNA2 | -2.139 | 0.0070 |
| MIR193A | -2.117 | 0.0102 |
| PRSS36 | -2.087 | 0.0090 |
| MAST1 | -2.076 | 0.0098 |
| CEBPA-DT | -2.056 | 0.0113 |
| TNFRSF14 | -2.055 | 0.0106 |
| ADAMTSL2 | -2.046 | 0.0127 |
| NPW | -2.041 | 0.0122 |
| CIMAP2 | -2.024 | 0.0144 |

We then examined the effects of another modified Ang II that can be produced through oxidant-mediated amino acid conversion. For these experiments, the same type of cells was treated with Ang II in which the Arg at position 2 was replaced with Pro (Arg2Pro), as in wild-type Ang II or Ang II (Arg2Glu). We found that Ang II (Arg2Pro) dramatically altered gene expression patterns (Fig. 6C). Compared to the dH_2_O-treated control, 2,473 genes were upregulated and 5,006 genes were downregulated by Ang II (Arg2Pro). Tables 7 and 8 list the names of the top twenty genes (in order of the extent of changes) that were modulated (upregulated and downregulated, respectively) by Ang II (Arg2Pro) vs. dH_2_O. Among the 45 genes upregulated by wild-type Ang II, 27 genes were also upregulated by Ang II (Arg2Pro). These genes are listed in Table 9. Genes that were upregulated by Ang II (Arg2Pro), but not by wild-type Ang II, include prostaglandin F receptor (PTGFR), which showed a 3-fold increase (calculated from log_2_ fold change of 1.632). Tables 10 and 11 list the names of the top twenty genes (in the order of the extent of changes) that were differentially regulated (upregulated and downregulated, respectively) by Ang II (Arg2Pro) relative to Ang II (wild-type). Thus, wild-type Ang II and Ang (Arg2Pro) differentially modify gene expression (Fig. 6E), indicating that the oxidant-mediated Arg-to-Pro conversion process substantially affects cell signaling elicited by Ang II.

**Table 7.** Genes upregulated in response to Ang II (Arg2Pro) treatment of cells relative to dH_2_O <u>control. Top 20 genes with highest fold change are listed.</u>

| Gene Name | Log<sub>2</sub> fold change | p-Value |
| --- | --- | --- |
| LINC02052 | 2.531 | 1.87 × 10<sup>-5</sup> |
| LINC01473 | 2.359 | 6.64 × 10<sup>-5</sup> |
| CAPS2-AS1 | 2.260 | 6.53 × 10<sup>-5</sup> |
| TET1P1 | 2.173 | 0.0002 |
| RGS2-AS1 | 2.106 | 0.0003 |
| LINC02726 | 2.071 | 0.0005 |
| PTGFR | 1.987 | 0.0004 |
| PWWP3B | 1.969 | 1.87 × 10<sup>-10</sup> |
| LVRN | 1.958 | 0.00071 |
| SPOPL-DT | 1.954 | 1.85 × 10<sup>-6</sup> |
| LRFN5-DT | 1.938 | 6.40 × 10<sup>-6</sup> |
| TTPA | 1.929 | 0.0011 |
| NHIP | 1.896 | 0.0005 |
| RFX8 | 1.873 | 0.0003 |
| IL18R1 | 1.865 | 0.0004 |
| LNCSRLR | 1.860 | 9.98 × 10<sup>-7</sup> |
| ALDH8A1 | 1.823 | 0.0013 |
| ANO3 | 1.816 | 7.98 × 10<sup>-6</sup> |
| RPS6KA6 | 1.812 | 1.19 × 10<sup>-10</sup> |
| LINC01762 | 1.812 | 0.0019 |

**Table 8.** Genes downregulated in response to Ang II (Arg2Pro) treatment of cells relative to <u>dH_2_O control. Top 20 genes with highest fold change are listed.</u>

| Gene Name | Log<sub>2</sub> fold change | p-Value |
| --- | --- | --- |
| CASZ1 | -2.608 | 1.22 × 10<sup>-6</sup> |
| DOK7 | -2.558 | 4.30 × 10<sup>-6</sup> |
| SLCO4A1 | -2.483 | 7.93 × 10<sup>-6</sup> |
| MIR10392 | -2.352 | 5.64 × 10<sup>-9</sup> |
| MAPK8IP1P1 | -2.345 | 1.32 × 10<sup>-8</sup> |
| PDZK1IP1 | -2.313 | 9.14 × 10<sup>-5</sup> |
| STRA6 | -2.233 | 3.87 × 10<sup>-5</sup> |
| TEX19 | -2.230 | 0.0002 |
| PITX1 | -2.228 | 1.03 × 10<sup>-6</sup> |
| SOD3 | -2.222 | 6.01 × 10<sup>-9</sup> |
| TMEM121 | -2.110 | 4.89 × 10<sup>-7</sup> |
| POMC | -2.100 | 9.63 × 10<sup>-5</sup> |
| CEND1 | -2.100 | 1.44 × 10<sup>-8</sup> |
| ESPNL | -2.093 | 1.55 × 10<sup>-6</sup> |
| SSTR3 | -2.075 | 0.0003 |
| ZNHIT2 | -2.056 | 6.51 × 10<sup>-12</sup> |
| SCNN1D | -2.049 | 4.16 × 10<sup>-5</sup> |
| RAB3IL1 | -2.048 | 3.25 × 10<sup>-8</sup> |
| CDHR1 | -2.044 | 0.0002 |
| BRSK2 | -2.034 | 0.0005 |

**Table 9.** List of genes commonly upregulated in response to the treatment of cells with Ang II (wild-type) and the treatment of cells with Ang II (Arg2Pro) relative to dH_2_O. Log_2_ fold change values for Ang II (wild-type) vs. log_2_ fold change values for Ang II (Arg2Pro) are also indicated.

| IGIP | STAG2 | RC3H1 |
| --- | --- | --- |
| 1.559 vs. 1.345 | 1.120 vs. 1.397 | 1.035 vs. 1.073 |
| PAK1IP1 | TMTC3 | ZBTB6 |
| 1.355 vs. 1.229 | 1.108 vs. 1.303 | 1.034 vs. 1.146 |
| CEP290 | NECAB1 | NMD3 |
| 1.246 vs. 1.395 | 1.098 vs. 1.038 | 1.023 vs. 1.232 |
| CERT1 | LRRC39 | RUFY3 |
| 1.188 vs. 1.044 | 1.095 vs. 1.357 | 1.013 vs. 1.159 |
| LRRC40 | SDHAF3 | LSM8 |
| 1.177 vs. 1.249 | 1.089 vs. 1.029 | 1.009 vs. 1.180 |
| ARL13B | THAP9-AS1 | PHAX |
| 1.175 vs. 1.104 | 1.077 vs. 1.377 | 1.001 vs. 1.107 |
| SMIM10L1 | CAVIN2 | HIF1A-AS2 |
| 1.170 vs. 1.267 | 1.062 vs. 1.258 | 1.000 vs. 1.182 |
| PHTF2 | IL18R1 | TNFAIP6 |
| 1.164 vs. 1.122 | 1.056 vs. 1.865 | 1.077 vs. 1.692 |
| ZBTB11 | SFR1P1 |  |
| 1.154 vs. 1.241 | 1.037 vs. 1.779 |  |
| CARNMT1-AS1 | SMCHD1 |  |
| 1.126 vs. 1.405 | 1.036 vs. 1.385 |  |

**Table 10.**
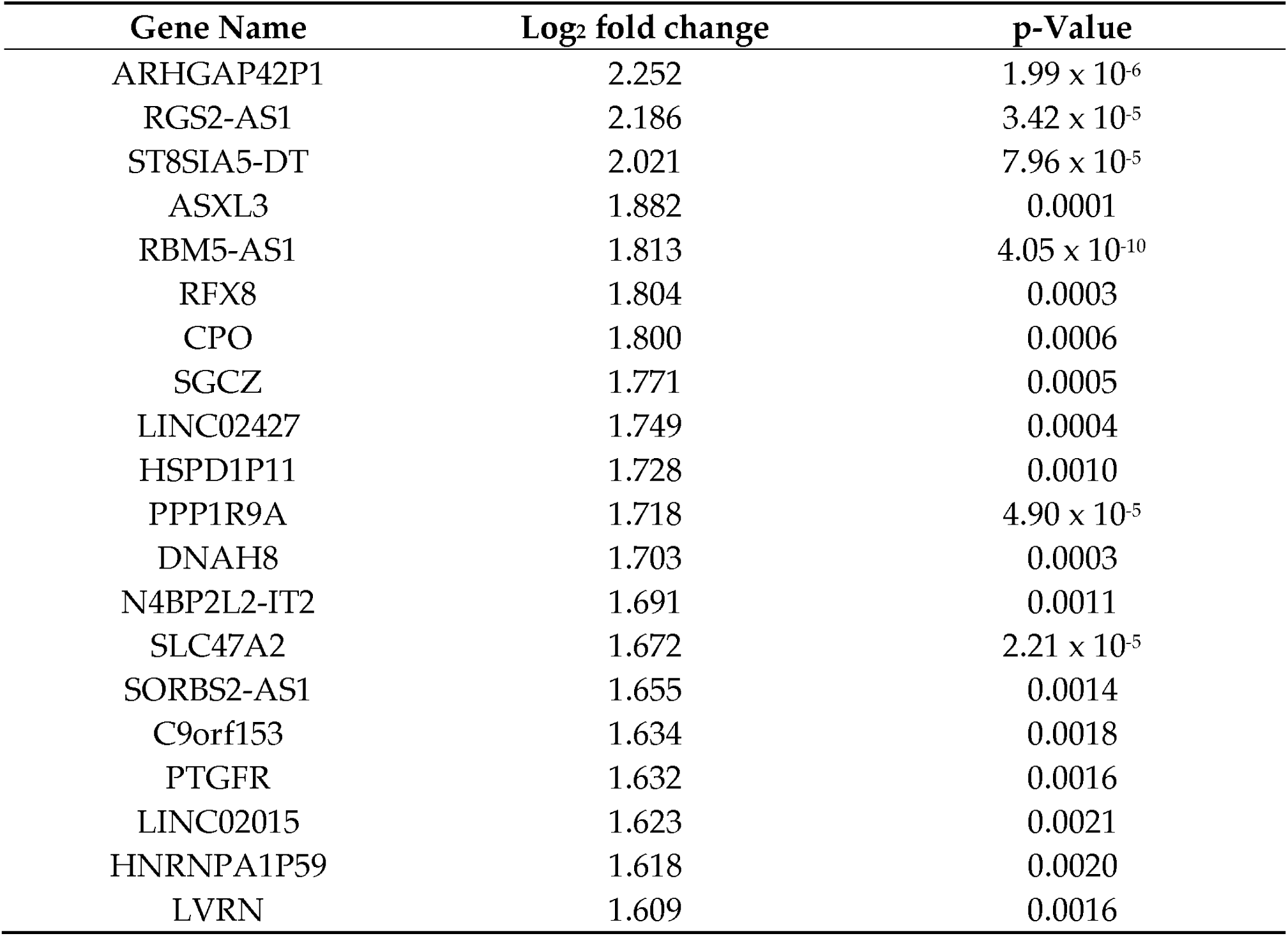
Genes upregulated in response to Ang II (Arg2Pro) treatment of cells relative to Ang <u>II (wild-type). Top 20 genes with highest fold change are listed.</u>

| Gene Name | Log<sub>2</sub> fold change | p-Value |
| --- | --- | --- |
| ARHGAP42P1 | 2.252 | $1.99 \times 10^{-6}$ |
| RGS2-AS1 | 2.186 | $3.42 \times 10^{-5}$ |
| ST8SIA5-DT | 2.021 | $7.96 \times 10^{-5}$ |
| ASXL3 | 1.882 | 0.0001 |
| RBM5-AS1 | 1.813 | $4.05 \times 10^{-10}$ |
| RFX8 | 1.804 | 0.0003 |
| CPO | 1.800 | 0.0006 |
| SGCZ | 1.771 | 0.0005 |
| LINC02427 | 1.749 | 0.0004 |
| HSPD1P11 | 1.728 | 0.0010 |
| PPP1R9A | 1.718 | $4.90 \times 10^{-5}$ |
| DNAH8 | 1.703 | 0.0003 |
| N4BP2L2-IT2 | 1.691 | 0.0011 |
| SLC47A2 | 1.672 | $2.21 \times 10^{-5}$ |
| SORBS2-AS1 | 1.655 | 0.0014 |
| C9orf153 | 1.634 | 0.0018 |
| PTGFR | 1.632 | 0.0016 |
| LINC02015 | 1.623 | 0.0021 |
| HNRNPA1P59 | 1.618 | 0.0020 |
| LVRN | 1.609 | 0.0016 |

**Table 11.** Genes downregulated in response to Ang II (Arg2Pro) treatment of cells relative to Ang II (wild-type). Top 20 genes with highest fold change are listed.

| Gene Name | Log<sub>2</sub> fold change | p-Value |
| --- | --- | --- |
| RRAD | -2.129 | $2.75 \times 10^{-14}$ |
| TMEM121 | -2.053 | $1.63 \times 10^{-13}$ |
| PITX1 | -2.042 | $4.23 \times 10^{-5}$ |
| PYCR3 | -2.030 | $4.56 \times 10^{-20}$ |
| SPATA2L | -1.957 | $3.46 \times 10^{-13}$ |
| ZNHIT2 | -1.953 | $8.41 \times 10^{-17}$ |
| NUDT8 | -1.943 | $6.44 \times 10^{-9}$ |
| F12 | -1.941 | 0.0001 |
| TSPO | -1.906 | $7.05 \times 10^{-18}$ |
| TRAPPC5 | -1.901 | $5.01 \times 10^{-16}$ |
| SOD3 | -1.898 | $2.33 \times 10^{-9}$ |
| STRA6 | -1.898 | 0.0002 |
| ATP5F1D | -1.897 | $4.10 \times 10^{-14}$ |
| SEPTIN1 | -1.896 | $3.24 \times 10^{-7}$ |
| DOK7 | -1.891 | 0.0003 |
| PRSS36 | -1.890 | $1.46 \times 10^{-6}$ |
| TECRP1 | -1.889 | $1.63 \times 10^{-19}$ |
| RNPEPL1 | -1.876 | $1.62 \times 10^{-17}$ |
| SLC39A4 | -1.871 | $1.64 \times 10^{-10}$ |
| PRR7 | -1.867 | $2.05 \times 10^{-14}$ |

## Discussion

The present study demonstrated that Arg2 of Ang II plays an important role in the interactions between Ang II and ACE2 that underlie its catalytic activity. Our experiments point to the importance of positively charged amino acids, as replacing Arg with the negatively charged Glu eliminated Ang II’s ability to compete with the fluorogenic substrate. We also found that Ang I and Ang III can enter the catalytic substrate tunnel and compete with the fluorogenic substrate (Mca-Ala-Pro-Lys-Dnp), and Arg plays a crucial role. Our experimental results, along with those from molecular dynamics simulations, provided mechanistic insights into the enzymology of ACE2.

As we used angiotensin peptides with Arg at the second position, which were replaced with other amino acids, these exhibited differential effects on ACE2 relative to wild-type peptides. The present study also found that some Arg2-modified angiotensin peptides occur naturally and exhibit biologic functions. This is based on our previously proposed hypothesis that oxidation can cause post-translational conversions of amino acid residues in biological systems [Suzuki & Hao, 2017; Suzuki, 2019; Suzuki 2022; Bablu et al., 2026].

Oxidative stress results from the production of reactive oxygen species (ROS), which can damage DNA, proteins, lipids, and small molecules [Halliwell & Gutteridge, 2015; Pizzino et al., 2017]. Protein carbonylation is a form of protein oxidation implicated in aging and various diseases [Levine, 2002; Dalle-Donne et al., 2003]. Primary protein carbonylation involves the direct oxidation of amino acid side chains [Suzuki, 2010]. We previously discovered a role of primary protein carbonylation in ligand/receptor-mediated cell signaling [Wong et al., 2008]. In these studies, we observed that the kinetics of ligand-mediated protein carbonylation are transient. Typically, in cultured cells, ligands induce protein carbonylation within 10 min, and protein carbonyls return to baseline by 30 min. These results led to our discovery of a protein de-carbonylation process [Wong et al., 2008]. To understand the mechanism of protein de-carbonylation, we tested the hypothesis that protein carbonyls could be reduced. We found that adding thiol reductants to tissue homogenates decreased protein carbonyl content [Wong et al., 2013]. In contrast, these thiol reductants had no effects on the carbonyl content of purified proteins, suggesting that protein carbonyls are reduced by components present in the biological system. Based on these results, we proposed that cells possess catalysts capable of reducing protein carbonyls. This concept is supported by our results demonstrating that heating tissue homogenates to inactivate cellular enzymes or knocking down glutaredoxin 1 in cells with siRNA inhibited protein de-carbonylation [Wong et al., 2013]. Thus, the biological system may contain catalysts that reverse the proline-to-glutamic semialdehyde reaction.

Since the oxidation of both Arg and Pro residues results in the formation of the same product, glutamic semialdehyde, Arg residues could be converted to Pro within the protein structure if the glutamic semialdehyde is reduced back to Pro. Moreover, the glutamic semialdehyde can be further oxidized to glutamic acid (Glu). Thus, it was proposed that, through glutamic semialdehyde, Arg, Pro and Glu could occur interchangeably in the protein structure, resulting in outcomes achieved by protein engineering or site-directed mutagenesis, but in this case, post-translationally and naturally [Suzuki, 2019].

Since Ang II contains Arg at position 2, we hypothesized that Arg-to-Glu and Arg-to-Pro conversions through the formation of glutamic semialdehyde could occur in this peptide molecule in the biological system during various oxidative stress conditions [Bablu et al., 2026]. The present study indeed demonstrated using mass spectrometry the occurrence of Arg-modified Ang I, Ang II, and Ang III in the biological system. To assess the significance of the modified peptides, we synthesized Arg2Glu- and Arg2Pro-converted Ang II peptides and examined their effects on gene expression in vascular smooth muscle cells using next-generation RNA-Seq. We demonstrated that each of these two modified peptides exerts differential effects on gene expression compared to wild-type Ang II.

Based on the Central Dogma of Molecular Biology, it has generally been thought that protein sequences are defined by nucleic acid sequences. In fact, in the field of protein engineering, site-directed mutagenesis changes DNA sequences to generate proteins with altered amino acid sequences. However, the proposal of oxidant-mediated protein amino acid conversion [Suzuki & Hao, 2017; Suzuki, 2019] challenged this concept by suggesting that Arg, Pro, and Glu residues in protein structures can be interchangeable. This is because the ROS-mediated protein carbonylation process produces a glutamic semialdehyde from both Arg and Pro residues, which can be further oxidized to Glu. Therefore, the conversions of amino acid residues that can be achieved by protein engineering/site-directed mutagenesis could also occur naturally and post-translationally through redox reactions.

The present study extended this concept to amino acid conversions that may occur in smaller peptides. Since a highly clinically important pathophysiologic mediator, Ang II, contains an Arg residue, we synthesized Ang II peptides with Arg at Position 2 replaced with either Glu or Pro and tested their ability to alter gene expression patterns in cultured vascular smooth muscle cells using RNA-Seq, as compared to the effects of wild-type Ang II.

One notable gene whose expression was increased by Ang II (Arg2Glu) but not by wild-type Ang II is FGF10, which has been shown to play protective roles against pulmonary vascular defects [Hadzic et al., 2023; Chao et al., 2019]. This suggests that this oxidized Ang II may counter the pathogenic actions of wild-type Ang II [Nádasy et al., 2024], perhaps through a different receptor.

Unlike wild-type Ang II, Ang II (Arg2Glu) downregulates multiple genes. In the context of cell signaling, it is difficult to assess the mechanisms underlying these downregulation events, as the signal transduction leading to gene downregulation is not well studied. Our laboratory previously reported a ligand-mediated dephosphorylation signaling mechanism regulated by endocytosis [Shults et al., 2018], and such signaling mechanisms may be elicited by oxidized Ang II to downregulate genes. One gene identified as downregulated by Ang II (Arg2Glu) is phosphodiesterase 6G (PDE6G). While PDE6G has not been recognized in smooth muscle cells [Cote et al., 2022], the present study identified the mRNA expression of this gene, opening a new concept regarding the role of PDE6G in smooth muscle biology. Downregulation of this gene by Ang II (Arg2Glu) may indicate that this oxidized Ang II may regulate smooth muscle contraction and growth in a manner distinct from that of wild-type Ang II.

In contrast to Ang II (wild-type), which modulated 45 genes under the experimental conditions and analytic parameters we used, Ang II (Arg2Pro) modulated 7,479 genes. Among 2,473 genes upregulated by Ang II (Arg2Pro), one notable gene is PTGFR, which regulates blood pressure [Zhang et al., 2010]. Another Ang II (Arg2Pro)-upregulated gene ARHGAP1 (Rho GTPase-activating protein 1), has been shown to regulate NADPH oxidase and superoxide production [Lőrincz et al., 2014]. One of the Ang II (Arg2Pro)-downregulated genes, SOD3 (extracellular superoxide dismutase), also regulates superoxide levels [Sasaki et al., 2021].

Given that Ang II oxidation occurs in the extracellular space, changes in the gene expression of this extracellular antioxidant may regulate oxidized Ang II levels. While the Arg-to-Glu conversion should occur through oxidation in the extracellular space, the Arg-to-Pro conversion also requires reduction reactions to convert glutamic semialdehyde to proline. We previously provided experimental evidence that Arg-to-Pro conversion indeed occurs on peroxiredoxin 6 protein [Suzuki, 2022], but this process likely occurs intracellularly. In the case of Ang II, it would be interesting to identify extracellular reductive processes that can drive this conversion.

In summary, Arg2 of Ang II plays an important role in ACE2 substrate interactions. Arg can be converted to Glu or Pro, and the present study identified novel angiotensin peptides in which the Arg residue is replaced with Glu or Pro that occur naturally in biological systems. Our confirmation that Ang II (Arg2Glu) and Ang II (Arg2Pro) exhibit biological actions distinct from those of wild-type Ang II has opened up the way for future research to determine whether these oxidatively-modified peptides indeed serve as mechanisms of Ang II action in various pathophysiological events.

